# Towards site-specific information on PET degrading enzymes using NMR near operational temperature

**DOI:** 10.1101/2024.09.10.612188

**Authors:** Valeria Gabrielli, Jelena Grga, Sabine Gavalda, Laura Perrot, François-Xavier Cantrelle, Emmanuelle Boll, Guy Lippens, Cyril Charlier

## Abstract

PETases are enzymes that can break down the poly-ethylene terephthalate (PET) polymer in its constituent building blocks. This enzymatic recycling process offers a sustainable solution for producing new, high-quality plastics from previously used materials. NMR spectroscopy can help in understanding and ultimately improving these enzymes but is always confronted with the lengthy step of acquisition and interpretation of triple resonance spectra for the spectral assignment. Here, we explore whether this step can be made more efficient by recording the spectra directly at high temperature, which simultaneously corresponds to more realistic working conditions for the enzyme. Taking the inactive variant of LCC^ICCG^ as an example, we compare spectral quality at 30°C and 50°C, and find that the latter condition greatly improves the Signal-to-Noise (S/N) ratio of the standard triple resonance spectra. Going up to 60ºC, we show that pulse sequences mainly used for the assignment of intrinsically disordered proteins (IDPs) also become feasible. As a result, we present a methodology enabling exhaustive backbone assignment based on a minimal set of triple resonance spectra acquired and analysed in less than two weeks. The assignment process hence can be completed on a time scale comparable to crystallography, bringing NMR in a favourable position to contribute to bio-structural studies on this family of highly thermostable PETases.

## 1. Introduction

Protein Nuclear Magnetic Resonance (NMR) in the field of structural biology is often perceived as laborious and limited to laboratories expert in spectroscopy. Indeed, every step, from sample preparation (including stable isotope labeling) to data acquisition (with multiple triple resonance experiments) and data analysis (with notably the resonance assignment) is time consuming, expensive and non-trivial, thereby restraining its use to specialized NMR facilities. Additionally, the assigned ^1^H-^15^N correlation spectrum that results from these efforts is only the starting point for more interesting insights about interactions, dynamics, *etc*. Important efforts are ongoing to improve all individual steps of the process with cell-free sample preparation for labeling (Hoffmann et al., 2018), novel interfaces facilitating the data acquisition for the non-specialist (Favier and Brutscher, 2019; Vallet et al., 2020) or improved resonance assignment procedures with automated and robust computational approaches (Klukowski et al., 2023; Pritišanac et al., 2019). However, NMR studies of proteins of molecular weight of ∼25 kDa and above continue to struggle with the necessity to acquire and analyse a large set of sophisticated tri-(and even higher) dimensional experiments. Although the use of methods to speed up data acquisition (Non-Uniform Sampling (Hoch et al., 2012), SO-FAST (Schanda et al., 2005)) can be very useful to further decrease the (precious and expensive) experimental time, they do not lower the barrier to its use as they often require further tweaking by experienced NMR spectroscopists. Contemporary NMR spectroscopy clearly needs to take the best of these advancements to open up to new horizons.

In the field of biotechnology, enzyme optimization is for many programs the key towards the success of bio-based approaches (Katsimpouras and Stephanopoulos, 2021). Computational methods based on artificial intelligence (Kouba et al., 2023; Reisenbauer et al., 2024; Yang et al., 2024) thereby can generate huge libraries of potential candidates, and coupled to high-throughput screening methods (Bozkurt et al., 2025; Yew et al., 2025), have the potential to rapidly converge upon improved enzymes. If NMR spectroscopy wants to participate in this effort – and can convince that its site-specific information is most useful for example to restrict the list of potential mutation sites for directed evolution (Bhattacharya et al., 2022; Gutierrez-Rus et al., 2024), obtaining the fully assigned ^1^H-^15^N spectrum as rapidly as possible seems a strict requirement.

One recent example where enzymes are at the core of a promising industrial process is bio-enzymatic recycling of plastic (Qiu et al., 2024; Tournier et al., 2023). PET hydrolases (PETases) break the ester linkages in polyethylene terephthalate (PET), a common plastic used in the manufacturing of beverage bottles, food containers or clothing fibers. By breaking down PET into its constituent monomers, terephthalate and ethylene glycol, they offer an attractive avenue for recovery of the building blocks that are the starting material for pristine plastic with the same characteristics as the original raw material (Tournier et al., 2023). In this rapidly evolving field, many newly identified PETase enzymes from diversity (Buchholz et al., 2022; Lin et al., 2025; Seo et al., 2024) have been screened and used as starting points for further engineering. It proved essential to work close to the PET glass temperature of ∼70°C, where the plastic becomes more dynamic at the molecular level without yet crystallizing (Thomsen et al., 2022, 2023). The temperature of denaturation of the enzymes thereby has become an important factor next to their intrinsic catalytical efficacy. The initially mesophilic *Ideonella sakaiensis* (*Is*) PETase (Yoshida et al., 2016) with its T_m_ of 46.1°C was evolved to become the ThermoPETase (Tm 58.6°C; Son et al., 2019), FastPETase (Tm 56.9°C; Lu et al., 2022), DepoPETase (Tm 69.4°C ; Shi et al., 2023), DuraPETase (Tm 78.7°C ; Cui et al., 2021) and finally HotPETase (Tm 82.5°C; Bell et al., 2022). Derived from the thermophilic Leaf-branch Compost Cutinase (LCC) (Sulaiman et al., 2012), a thermostable cutinase characteristic from the type I group (Joo et al., 2018), its quadruple mutant LCC^ICCG^ was equally optimized based on its activity as well as its thermostability (*T*_*m*_ = 85.8ºC for LCC and 93.3ºC for LCC^ICCG^ at pH 8, both proteins without C-terminal His-tag) (Tournier et al., 2023). The latter enzyme has shown great promise for industrial application of plastic recycling (Arnal et al., 2023; Tournier et al., 2020).

From an NMR standpoint, this high thermostability could be - at least in theory-a significant advantage, as spectral recording at higher temperatures should lead to faster tumbling of the enzyme and hence mimic a lower molecular weight species. Here, taking the catalytically inactive S165A variant of LCC^ICCG^ as an example, we explore the possibility to facilitate the resonance assignment of the thermophilic PETases by recording a set of 3D experiments directly at higher temperatures. We first measure experimentally the global tumbling time (*τ*_c_) at different temperatures, and find a twofold reduction already when going from 30°C to 50°C. When comparing spectra at 30°C, temperature where we obtained our first assignments (Charlier et al., 2022), and similar experiments recorded at 50°C, we find that the standard triple resonance experiments work significantly better at 50°C, with an important gain in Signal-to-Noise (S/N) ratio and hence in spectral recording time. With the assignments at 50°C in hand, a temperature series of simple 2D spectra gives access to the assignments at both lower (30°C) and higher (68ºC), whereby the latter temperature corresponds to the operational industrial conditions (Arnal et al., 2023). We equally find that additional experiments based on the hNcaNNH and HncaNNH pulse sequences, mostly used for assignment of highly flexible Intrinsically Disordered Proteins (IDP) although equally applied with success to some larger enzymes after complete deuteration (Frueh et al., 2006; Frueh, 2014), become feasible without any deuteration, and this despite the 30 kDa MW range of the enzymes. Based on this observation we achieve *de novo* backbone assignment of an active variant of LCC^ICCG^) primarily based on the resulting ^15^N and ^1^H connectivities. Within 6 days of data acquisition followed by 3 days of data analysis, nearly 75% of the ^15^N/^1^H^N^ resonances were assigned. Such achievement should enable NMR to follow protein design at a pace at least comparable to X-ray crystallography and alleviate the bottleneck of resonance assignment in large scale projects involving thermostable enzymes.

## 2 Materials and Methods

### 2.1 Protein expression and purification

LCC^ICCG^-S165A and its active variant were expressed and purified according to previous studies (Charlier et al., 2022, 2024).

### 2.2 NMR spectroscopy

All details about NMR spectroscopy are given in the supporting information.

## 3 Results

### 3.1 Determination of *τ*_c_ as function of temperature

The global tumbling time *τ*_c_ (or, strictly speaking, the correlation time of the dipolar autocorrelation function of a given amide H-N vector (Cavanagh et al., 2007)) is one criterium that determines the linewidth of the corresponding amide correlation peak. Other factors such as chemical exchange of the amide protons with water or the presence of dynamics in the intermediate time scale regime can equally contribute. Whereas the ratio of transverse (T_2_) to longitudinal (T_1_) relaxation times can be used to estimate this *τ*_c_ parameter (Tjandra et al., 1995), the increasing influence of intramolecular motions on the T_1_ relaxation time for larger proteins (Luginbühl, 2002) has motivated the development of an alternative method. The TRACT (TRosy for rotAtional Correlation Times) method measures the decay of both TROSY and anti-TROSY components of the ^15^N doublet (Lee et al., 2006), and translates the differential rate into a value of *τ*_c_ without interference of dipole–dipole (DD) relaxation by remote protons or relaxation contributions from chemical exchange (Lee et al., 2006; Robson et al., 2021). TRACT experiments on the LCC^ICCG^-S165A enzyme were recorded at various temperatures ranging from 30ºC to 60ºC. By integrating the envelope of the proton spectra over the amide region as function of the relaxation delay, both relaxation rates were obtained using an exponential minimization (Figure 1. A & B). The difference between both rates (*ΔR*_2_) was interpreted using the equation given by Robson and coauthors (Robson et al., 2021) and yielded *τ*_c_ values varying from 13 to 5.7 ns upon increasing the temperature from 30°C to 60°C (Figure 1. C). In agreement with the values expected from the temperature dependence of water viscosity for a rigid body of this size (Garci a De La Torre et al., 2000) (Figure 1. D), this accelerated tumbling predicts a significant decrease in transverse relaxation rates for both amide and C*α* resonances (Yamazaki et al., 1994), thereby reducing the need for (partial) deuteration of the sample. We show here that because of this superior sensitivity, nearly complete assignment of the backbone and side chain spectra of LCC^ICCG^-S165A can be obtained with a minimal set of experiments recorded at 50°C (Mathematical details are given in Supplementary material).

**Figure 1.**
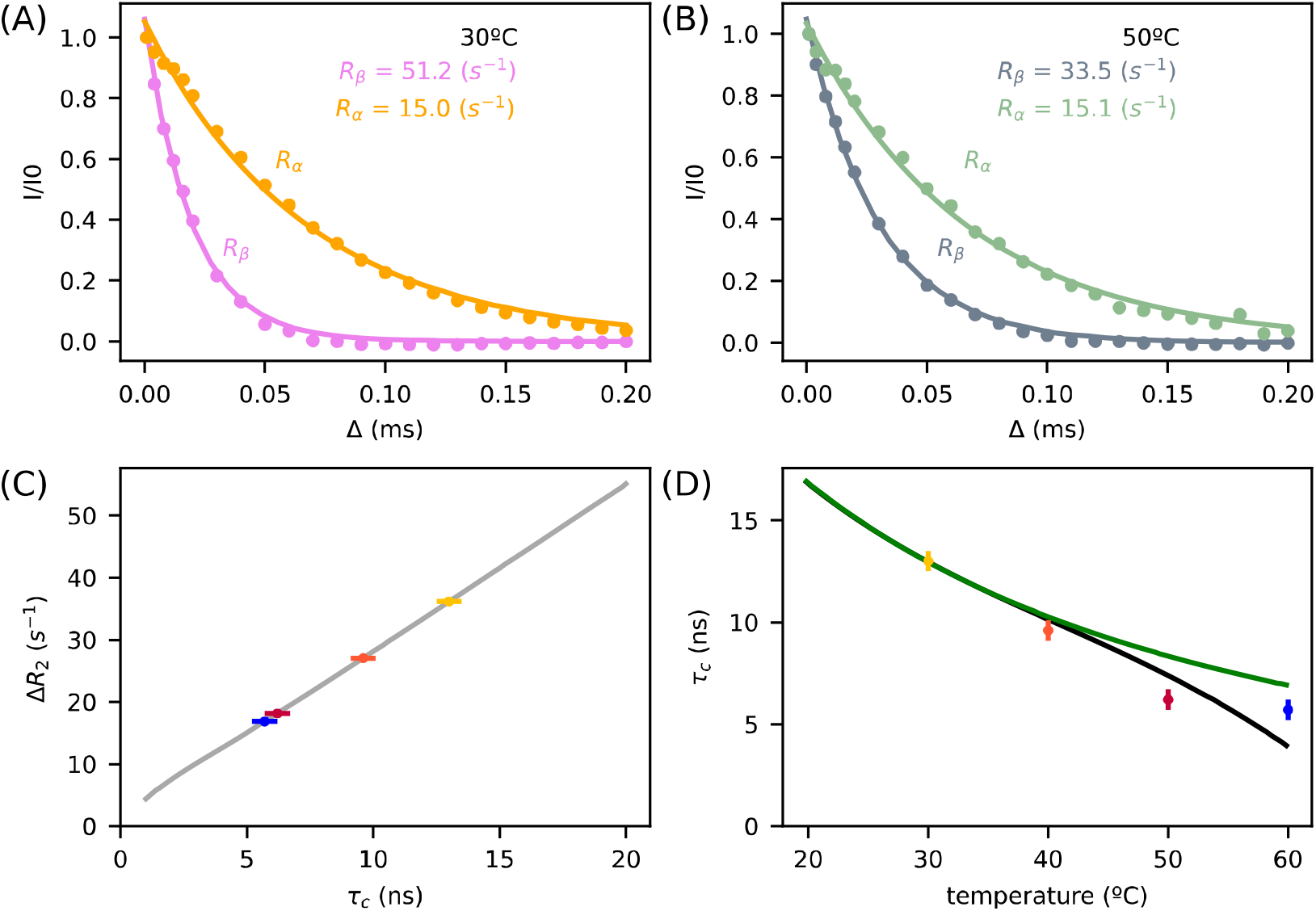
Determination of the correlation time *τ*_c_ as function of temperature. (**A-B**) TRACT analysis of LCC^ICCG^-S165A at (**A**) 30ºC and (**B**) 50ºC. Both panels display the single-spin-state ^15^N *R*_*α*_ and *R*_*β*_ relaxation decay curves obtained using the TRACT sequence. The dots represent the experimental points, and the lines represent the results of fitting the curves to a single exponential decay. (**C**) Theoretical curve of the difference between ^15^N relaxation rates, *ΔR*_2_ as function of *τ*_c_ following the equations of Robson and coworkers (Robson et al., 2021). (**D**) Theoretical curve of the temperature dependence of *τ*_c_ as predicted for a protein with its *τ*_c_ at 30°C fixed at 13 ns, based on the temperature dependence of water viscosity. Solid lines shows the *τ*_c_ values as a function of temperature as calculated with water viscosity according to Weast (Weast, 1979) (black) or Kestin *et al*. (Kestin et al., 1978) (green). On **C** and **D** panels, dots correspond to experimental values at 30ºC (yellow), 40ºC (orange), 50ºC (red) and 60ºC (blue). Error bars on (**C**) and (**D**) were calculated based on and estimated uncertainty of +/-0.5 s^-1^ for the fits of the *R*_*α*_ and *R*_*β*_ curves.

### 3.2 Influence of the temperature on spectral quality

To evaluate experimentally the influence of temperature on the spectral quality of our PETase, we recorded on the same sample at 30°C and 50°C both the ^1^H-^15^N HSQC and ^1^H-^15^N TROSY correlation spectra using standard Bruker pulse programs (*hsqcf3gpph19* and *trosyf3gpph19*). Already at 30ºC, signal intensity improved by ∼ 40% when comparing the HSQC experiment with the TROSY and going to 50ºC clearly boosted the intensity gain by another ∼15% (Figure 2.A). Equally important, even though cross peaks generally shift upfield with increasing temperature, (Asakura et al., 1995; Wagner and Wüthrich, 1979) the number of peaks in the 2D correlation spectra remained nearly the same between spectra recorded at both temperatures (Figure 2.B). Contrary to the case of IDPs, where amide proton signals at neutral to alkaline pH completely disappear at higher temperatures (Gil et al., 2013; Lopez et al., 2016), rapid amide proton exchange does not seem to represent a problem for this highly stable PETase.

**Figure 2:**
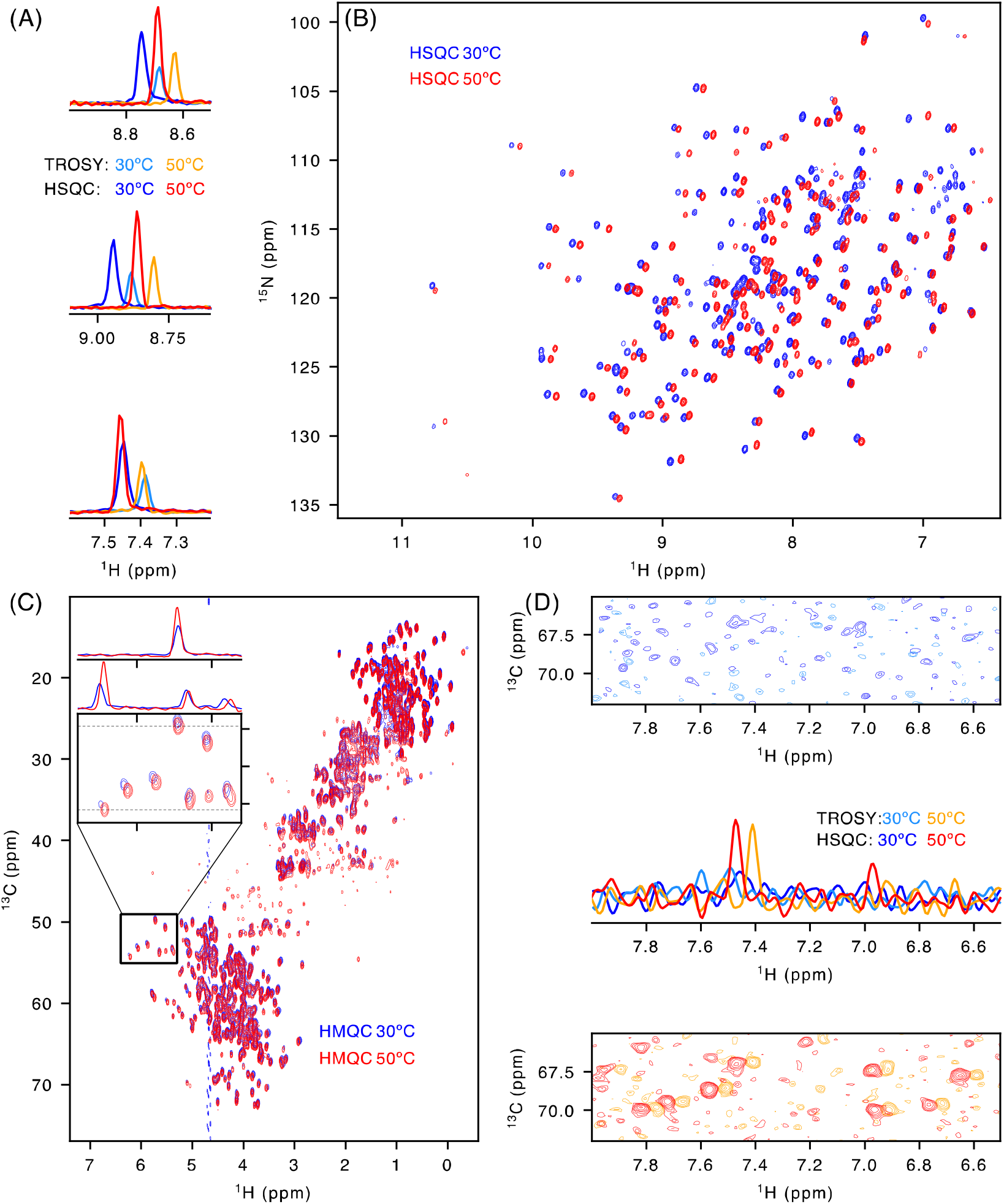
Selection of the best conditions and experiments for PETase assignment LCC^ICCG^-S165A. (**A**) 1D ^1^H slices through G88 (top), A59 (middle), G163 (bottom) cross peaks from ^1^H-^15^N HSQC at 30ºC (blue), ^1^H-^15^N TROSY at 30ºC (light-blue), ^1^H-^15^N HSQC at 50ºC (red) and ^1^H-^15^N TROSY at 50ºC (orange). (**B**) ^1^H-^15^N HSQC at 30ºC (blue) and 50ºC (red). Cross peaks generally shift upfield, but their number is the same at both temperatures (**C**) ^1^H-^13^C HMQC at 30ºC (blue) and 50ºC (red). The insets show how representative ^13^C*α* signals are sharper at 50ºC compared to 30ºC. The ^1^H 1D slices were taken at the ^13^C frequency displayed by the dashed lines. (**D**) Selected region of ^1^H-^13^C plane of HNCACB HSQC-based (blue) and TROSY-based (light-blue) at 30ºC (top) and of HNCACB HSQC-based (red) and TROSY-based (orange) at 50ºC (bottom). The central panel shows the 1D trace through the cross-peak at 67 ppm (^13^C).

In a typical backbone assignment experiment (*e.g*. HNCACB), the lengthy back-transfer of ^13^C*α* to ^15^N magnetization is often a limiting step due to the rapid transverse relaxation of the ^13^C*α* transverse magnetization. To evaluate how beneficial spectral recording at 50°C would be, we first measured ^1^H-^13^C HMQC spectra with a wide ^13^C spectral window both at 30ºC and 50ºC (Figure 2.C). Relative to the isolated methyl signal of M166 (Charlier et al., 2022), we observe a nearly 40% increase in intensity for ^1^H*α*-^13^C*α* resonances in the spectrum recorded at 50ºC compared to that at 30ºC. To evaluate the effect this has on triple resonance spectra, we acquired the ^1^H-^13^C planes of a HNCACB spectrum using the Bruker standard HSQC-based (*hncacbgpwg3d*) or TROSY-based (*trhncacbgp3d*) experiments both at 30ºC and 50ºC. Using a reasonable amount of spectrometer time (*e.g*. 8 scans per increment), the isolated ^13^C*β* signals of threonine residues hardly came out of the noise at 30ºC while they were clearly visible at 50ºC (Figure 2.D). The regular ^1^H-^15^N HSQC pulse sequence rather than its TROSY version being beneficial especially at 50°C (Figure 2. A & B), we tested both HNCACB sequences at 50°C. The advantage of the regular HSQC over the TROSY based HNCACB experiments persisted although less pronounced (Figure 2.D). Therefore, we recorded a HNCACB triple resonance spectrum without TROSY selection at 50ºC and pH 7.5. Starting from our previous assignment of the ^1^H-^15^N HSQC spectrum of LCC^ICCG^-S165A at 30ºC (Charlier et al., 2022), we used this single spectrum to confirm the assignments at 50°C (Figure S1). A HNCO experiment was recorded to obtain the ^13^C’ resonances at 50°C. Alltogether, 85% of the ^1^H^N^/^15^N resonances along with 91% for ^13^C*α* (234 out of 258), 90% ^13^C*β* (215 out of 239 residues, excluding the 19 glycines) and 85% of ^13^C’ resonances (203 out of 239, excluding all residues preceding a proline residue) were assigned.

### 3.3. Raising the temperature for hNcaNNH

In our general effort to accelerate the assignment of the ^1^H-^15^N spectrum, we explored the possibility to record a hNcaNNH experiment (Weisemann et al., 1993). While the common approach of backbone assignment consists of connecting consecutive residues through their various ^13^C resonances, this experiment together with related ones have been developed to obtain sequential correlations between successive amide proton and nitrogen resonances ^1^H_(i-1)_, ^15^N_(i-1)_, ^1^H_(i)_, ^15^N_(i)_, ^1^H_(i+1)_, ^15^N_(i+1)_ (Weisemann et al., 1993; Bracken et al., 1997; Sun et al., 2005). Although recognized as true game changers in the assignment process for disordered proteins (Lopez et al., 2013; Charlier et al., 2017), they have not been used for non-deuterated globular proteins of the size of PETases due to their inherent low sensitivity. Indeed, the bottleneck in these experiments is the fast ^13^C*α* relaxation rate that significantly deteriorates their sensitivity. To explore whether these experiments would profit from the same sensitivity gain as the above-described triple resonance spectra with a single period of ^13^C*α* transverse relaxation, we recorded a hNcaNNH directly at 50°C with the standard pulse sequence from the Bruker library (hncannhgp3d). The sensitivity of this experiment at 50ºC proved at the edge in a reasonable amount of experimental time (2 days 15 hours for a fully sampled dataset) (Figure 3.A). However, the signal-to-noise ratio further increased (1.5 to 2-fold) when raising the temperature to 60ºC (Figure 3.A) allowing indeed connection of consecutive residues through their ^15^N_(i-1)_ and ^15^N_(i+1)_ resonances (Figure 3.B).

**Figure 3.**
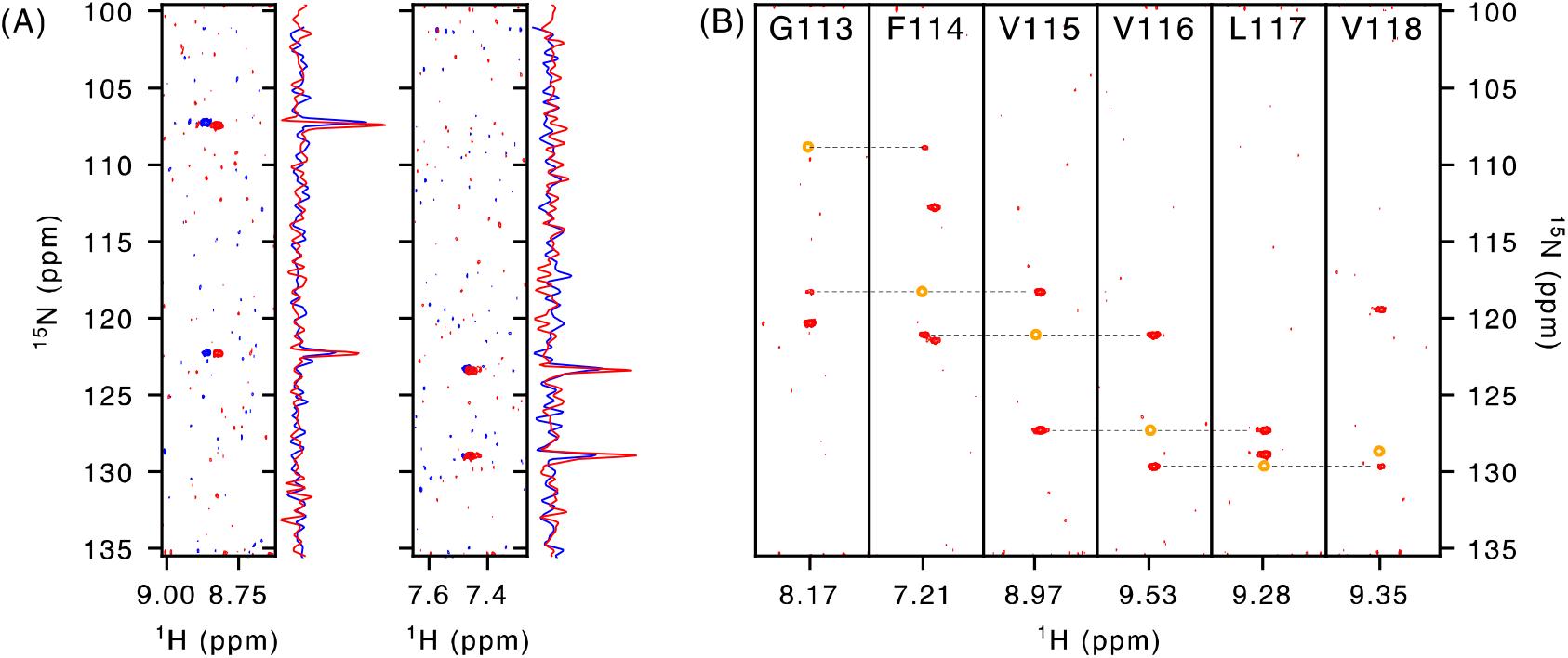
Connecting ^15^N resonances through hNcaNNH experiment. **(A)** Overlay of the ^1^H-^15^N strip extracted from the hNcaNNH at 50ºC (blue) and 60ºC (red). ^15^N traces were extracted at the ^1^H frequency of the selected signal. **(B)** ^1^H-^15^N strips of consecutive residues at 60ºC shown for residues from G113 to V118. Red resonances correspond to ^15^N(i-1) and ^15^N(i+1) while the autocorrelation cross-peak leading to the ^15^N(i) was added manually in orange based on the ^15^N value obtained from the ^1^H-^15^N HSQC spectra.

### 3.4 Rapid *de novo* backbone assignment

Based on the quality of the hNcaNNH acquired at 60ºC, we sought to investigate whether a *de novo* backbone assignment primarily based on the hNcaNNH and HncaNNH experiments would be possible. Turning stretches of connecting resonances into a backbone assignment further required a HNCACB spectrum to obtain amino acid type identifying ^13^C chemical shift values (Figure 4). Finally, we recorded HNCO and HNCACO spectra for independent verification of the obtained assignments. With a 500µM sample of the active enzyme in a 5mm tube, on a 900MHz spectrometer equipped with a triple-resonance cryogenically cooled probe head, the complete data set was acquired within 6 days of machine time (Table 1).

**Table 1.**
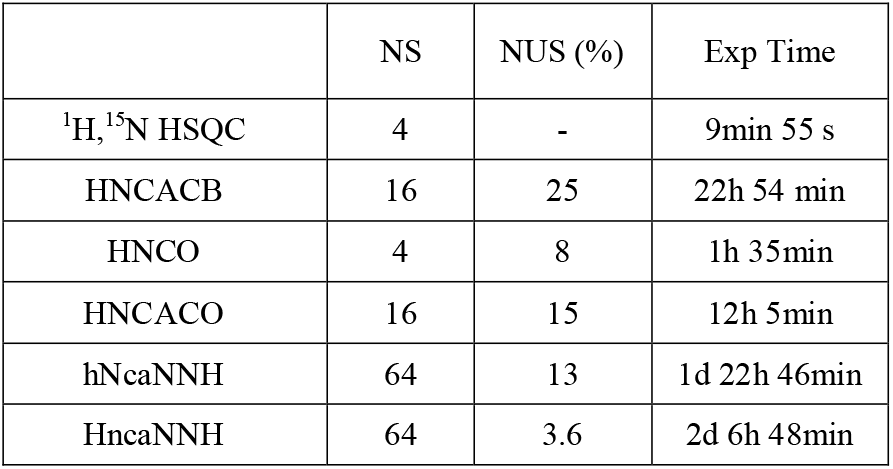
Summary of the experiments acquired within a week of machine time. More NMR parameters are given in the supplementary table.

**Figure 4.**
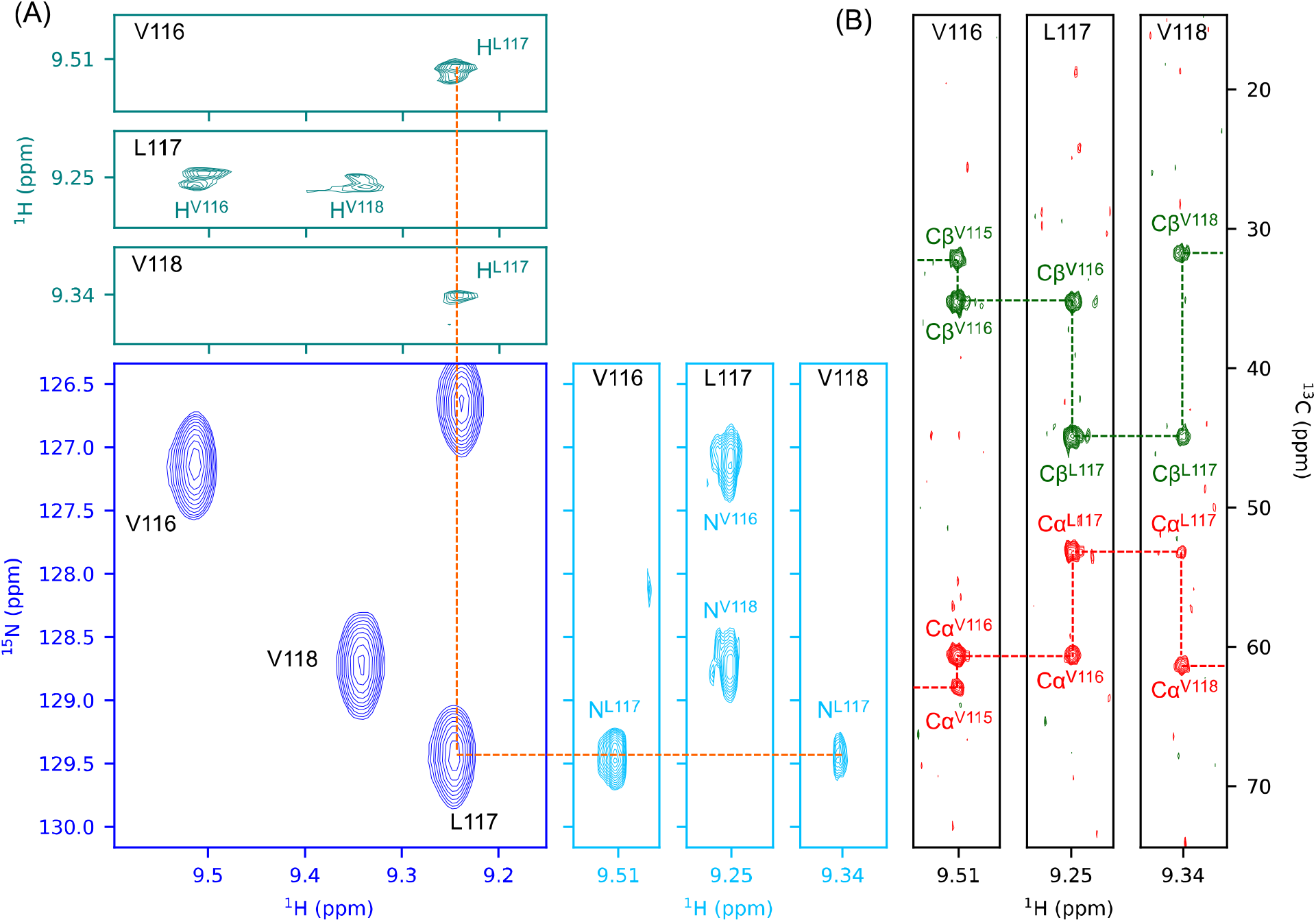
*De novo* assignment primarily based on ^15^N and ^1^Hconnectivities. **(A)** Example of connectivity between the HncaNNH (teal), HSQC (blue) and hNcaNNH (skyblue) experiments for residues V116-L117-V118. The ^1^H-^1^H and ^15^H-^1^H strips were extracted at the ^1^H^N^ resonance frequency of the 3 signals. The strips are only displayed in the window corresponding to the HSQC experiment (blue). The resonances of V115 and I119 respectively on the V116 and V118 strips are outside of displayed window. The orange line shows the connectivity for L117 both from V116 (i-1) and L118 (i+1). **(B)** HNCACB strips displayed for all 3 residues with the C*α* in red and C*β* in green.

The elevated temperature (60ºC) led to improved signal-to-noise data and further allowed efficient non-uniform sampling providing very high-resolution spectra. Especially for the HncaNNH experiment, this enhanced resolution proved important to establish firm connectivities between resonances of neighboring residues and thereby to speed up significantly the assignment procedure. In complement, HNCACB was used to establish the directionality of the assignment due its capacity to point i and i-1 carbon resonances. After 3 days of spectral analysis carried out by one of us (E.B.), initially external to the project and hence without any pre-knowledge of previous PETase assignments but with extensive experience in the use of CCP-NMR (Skinner et al., 2016), nearly 80% of backbone (^1^H/ ^15^N) resonances were assigned. The slightly lower percentage than the one obtained previously for the inactive LCC^ICCG^ variant is primarily due to additional broadening of some resonances assigned to solvent exposed residues that were already at the limit of observation at 50ºC in LCC^ICCG^-S165A (T211 and G127; see Figure S1).

## 4 Discussion

The initial steps of the development of new or improved PETases have invariably involved molecular modeling and/or bioinformatics, whereby the starting 3D structures that served as input for these efforts mostly came from X-ray crystallography. Examples such as *Tf*Cut2 (PDB code: 4CG1 (Roth et al., 2014)), Cut190 (5ZNO (Numoto et al., 2018)), LCC and mutants (4EBO (Sulaiman et al., 2012) and 6THS, 6THT (Tournier et al., 2020)) or *Is*PETase (5XG0 (Han et al., 2017)), are reviewed by Liu *et al*. (Liu et al., 2023) and Tournier *et al*. (Tournier et al., 2023).

Solution biomolecular NMR has only recently entered this field, and not as much as to obtain structural information but to characterize the interaction between the enzyme and small soluble substrates (Charlier et al., 2022; Hellesnes et al., 2023) or to investigate the role of the catalytic histidine with its associated pKa value (Charlier et al., 2024). Recently, a solid state NMR study of the *Tf*Cut2 enzyme embedded in a trehalose glass in the presence of nanoparticles (NPs) made of ^13^C labeled PET oligomers equally added to our understanding of the enzyme/polymer interaction and the biocatalytic mechanism of the hydrolase reaction (Falkenstein et al., 2023). However, if NMR wants to consolidate its place in this large-scale project that is the development of industrial PETases, easy, rapid and robust methods providing the assignment of the ^1^H-^15^N and/or ^1^H-^13^C spectra are required.

Particularly true for enzymes near or beyond 30kDa, considered the size-limit of classical solution state NMR, assignment of the ^1^H, ^15^N HSQC spectrum remains a real bottleneck for many projects in which NMR is or could be involved. Except for intrinsically disordered proteins (IDPs), where the relevant *τ*_c_ parameter does not correlate with the MW of the protein, the size problem leads not only to an increased number of peaks but also to peak broadening and hence decreased signal-to-noise. Deuteration (Gardner and Kay, 1998; LeMaster and Richards, 1988; Sattler and Fesik, 1996) together with TROSY approaches (Pervushin et al., 1997) can alleviate the latter peak broadening, but significantly increases the price per sample, with associated costs that become an important factor to consider in a program that generates many variants. Alltogether, we observe in the Biomagresbank (Hoch et al., 2023) an exponentially decreasing number of deposited assignments for proteins larger than 150 residues, and finally to no more than ∼ 100 entries for proteins of the size of LCC and its derivatives (Klukowski et al., 2023).

Although the NMR assignment clearly will never keep up with the many PETase variants that are currently tested in academic and industrial laboratories all over the world, speeding up the process could increase its use in this rapidly evolving field.

Our present study of LCC^ICCIG^ with all NMR experiments recorded directly on a simple sample at high temperature demonstrates that one can indeed significantly accelerate the process. Not only do we increase the sensitivity of the standard triple resonance experiments that are the cornerstone of the assignment process by matching ^13^C frequencies (Figure 2.D), but directly connecting consecutive amide peaks through the complementary hNcaNNH or HncaNNH spectra becomes possible (Figure 3, Figure 4) and dramatically speeds up the procedure. High temperatures equally imply we are studying these enzymes in what we would call ‘near-working industrial conditions” (Arnal et al., 2023) (to make the parallel with “near-physiological conditions” for disease related enzymes), and should therefore be seen as a win-win situation for solution state NMR : it not only yields higher-quality data and opens the door to experiments that would otherwise be impossible, but it also gives information about the enzyme in conditions that approximate its real-world functioning.

Based on the present study, we can fairly estimate that in less than three weeks, one could go from sample preparation (expression and purification of the isotopically labelled sample) to the obtention of an assigned ^1^H-^15^N spectrum. NMR thereby can become complementary to X-ray crystallography as a structural method to study PETases and potentially other thermophilic enzymes in solution.

## Authors’ contribution

V.G., E.B. and F.-X.C. recorded and analysed NMR spectra. J.G., L.P. and S.G. resources. G.L. and C.C. conceptualization, project administration, supervision, writing – original draft preparation, writing - review & editing. All authors reviewed the manuscript.

## Acknowledgements

We would like to thank Alain Marty, Vincent Tournier, Nicolas Chabot, and Marc Gueroult (Carbios, France) for insightful discussions and comments on the manuscript. We thank the ICEO facility of the Toulouse Biotechnology Institute (TBI), which is part of the Integrated Screening Platform of Toulouse (PICT, IBiSA), for providing access to protein-purification equipment. We thank Dr. E. Cahoreau and L. Peyriga in the MetaToul (Toulouse metabolomics & fluxomics facilities, www.metatoul.fr) NMR facility. MetaToul is part of the French National Infrastructure for Metabolomics and Fluxomics MetaboHUB-AR-11-INBS-0010 (www.metabohub.fr), and is supported by the Région Midi-Pyrénées, the ERDF, the SICOVAL and the French Minister of Education & Research, who are all gratefully acknowledged. Financial support from the IR INFRANALYTICS FR2054 for conducting the research is gratefully acknowledged. This study was supported by the OPTIZYME grant financed by ADEME - contract number 2282D0513-C.

## Data availability

All the parameter sets for the NMR spectra have been deposited to https://zenodo.org/records/15425383 and/or contact either

