## Supplementary Materials for "Towards site-specific information on PET degrading enzymes using NMR near operational temperature"

Valeria Gabrielli^1^, Jelena Grga^1^, Sabine Gavalda^2^, Laura Perrot^2^, François-Xavier Cantrelle^3,4^, Emmanuelle Boll^3,4^, Guy Lippens^1,*^, Cyril Charlier^1,*^

^1^Toulouse Biotechnology Institute (TBI), University of Toulouse, CNRS, INRAE, INSA Toulouse, 135 Avenue de Rangueil, 31077 Toulouse Cedex 04, France

^2^Carbios, Parc Cataroux – Bâtiment B80, 8 rue de la Grolière, 63100 Clermont-Ferrand, France

^3^CNRS EMR 9002 ─ Integrative Structural Biology, F-59000 Lille, France.

^4^Univ. Lille, Inserm, CHU Lille, Institut Pasteur de Lille UMR 1167 ─ RID-AGE ─ Risk Factors and Molecular Determinants of Aging-Related Diseases, F-59000 Lille, France.

### 1. **NMR Spectroscopy**

#### 1.1 LCC^ICCG^-S165A

All experiments recorded for backbone assignment at 50ºC were acquired on a 800 MHz spectrometer equipped with a 5-mm cryoprobe with z pulsed field gradients. The sample contained 580 μM of ^15^N-^13^C labeled protein in 25 mM Tris-HCl buffer pH 7.5 with 100 mM NaCl. The data presented here were recorded using the Bruker library pulse sequences: 2D ^1^H-^15^N HSQC, HNCACB and HNCO. All spectra were processed using Topspin 4.0.8 (Bruker Biospin). Spectral analysis for backbone assignments were performed manually using POKY (Lee et al., 2021). One important experimental factor we found when measuring at higher temperatures is the necessity to seal the sample. Otherwise, already after a day, evaporation set in and shim quality rapidly deteriorated. Spectral dimensions for the HNCO experiments were 3622 Hz (F_1_)ｘ2919 Hz (F_2_)ｘ12500 Hz (F_3_) corresponding to 18ｘ36ｘ15.6224 ppm, with sampling durations of 15.5 ms (t_1_), 10.9 ms (t_2_), 160 ms (t_3_). Spectra were centered at 4.700 ppm (^1^H), 117.5 ppm (^15^N), 173 ppm (^13^CO). Spectral settings for the HNCACB experiment were 6438.5 Hz (F_1_)ｘ2919 Hz (F_2_)ｘ14084.25 Hz (F_2_) corresponding to 32ｘ36ｘ50 ppm, with sampling durations of 14.9 ms (t_1_), 10.9 ms (t_2_), 0.009 ms (t_3_). Spectra were centered at carriers were set to 4.7 ppm (^1^H), 117.5 ppm (^15^N), 40 ppm (^13^Cα). Both experiments were acquired with 4 scans per increment.

#### 1.2 Active variant of LCC^ICCG^

Experiments were recorded at 60ºC on a sample containing 600 μM of ^15^N-^13^C labeled of the active variant of LCC in 25 mM Tris-HCl buffer pH 7.5 with 100 mM NaCl on a 900 MHz spectrometer equipped with a 5-mm cryoprobe with z pulsed field gradients with the following parameters:

|  | Time domain data size (points) | | | Spectral width/Carrier frequency (ppm) | | | NS | Delay time (s) | NUS (%) | Pulse program |
| --- | --- | --- | --- | --- | --- | --- | --- | --- | --- | --- |
|  | t1 | t2 | t3 | F1 (^1^H) | F2(^15^N) | F3 |  |  |  |  |
| ^1^H,^15^N HSQC | 3076 | 128 |  | 19.83/4.7 | 35/117 |  | 4 | 1 | - |  |
| HNCACB | 4000 | 128 | 128 | 15.42/4.7 | 35/117 | 60/42 (^13^C) | 16 | 1 | 25 | hncacbgpwg3d |
| HNCO | 4000 | 128 | 112 | 15.42/4.7 | 35/117 | 13/173.5 (^13^C) | 4 | 1 | 8 | hncogpwg3d |
| HNCACO | 4000 | 128 | 112 | 15.42/4.7 | 35/117 | 13/173.5 (^13^C) | 16 | 1 | 15 | hncacogpwg3d |
| (H)N(CA)NNH | 2048 | 128 | 128 | 13.88/4.7 | 35/117 | 35/117 (^15^N) | 64 | 1 | 13 | hncannhgp3d |
| H(NCA)NNH | 4000 | 128 | 512 | 15.42/4.7 | 35/117 | 6/8 (^1^H) | 64 | 1 | 3.6 | hncannhgp3d.2 |

NUS percentages were estimated based on the number of resonances expected in each dataset. Data were reconstructed using standard MDD software (Mayzel et al., 2014). Spectral analysis and backbone assignments were performed manually using CcpNmr analysis software v2.5 (Vranken et al., 2005).

### 2. **TRACT analysis**

TRACT spectra were recorded with the pulse sequence of Lee et al., implemented as individual experiments for the α and β ^15^N spin states (Lee et al., 2006). A relaxation delay of 3s was used between scans, and 4k points were recorded. A total of 24 delays ranging from 1 to 200 ms were sampled as increments in a pseudo-2D matrix, and recorded with 512 scans per delay. After manual phasing and base line correction, the integral of the [10.0-6.4] ppm region was determined and plotted as a function of the corresponding delay. The resulting curve was fitted as a mono-exponentially decaying function, and the difference between the rates of both components was interpreted in terms of τ_c_ as described by Robson et al (Robson et al., 2021).

Theoretical calculus of the τ_c_ value as a function of temperature was performed by fixing the τ_c_ value at the experimentally determined value of 13ns for 30°C, and then applying the following formula (Garcı́a De La Torre et al., 2000) :

$$\tau_{c}\left( 30ºC \right)=\frac{\eta_{30ºC}}{\eta_{T}}*\frac{273+T}{303}*\tau_{c}(T)$$

where η_T_	is the viscosity of water at the temperature T (using the Celsius scale). The latter was calculated using the equation from (Weast, 1979)

$$\eta_{T}=1.7753-0.0565T+1.0751*{10}^{-3}T^{2}-9.222*{10}^{-6}T^{3}$$

or, alternatively, from (Kestin et al., 1978):

$$log({\eta_{T}}/{\eta_{20})}=\frac{\left( 20-T \right)}{\left( T+96 \right)}\times1.2378-{1.303*10}^{-3}\left( 20-T \right)+{3.06*10}^{-6}\left( 20-T \right)^{2}+{2.55*10}^{-8}\left( 20-T \right)^{3}$$

and does not consider that our protein sample was not in pure water but rather in 25mM Tris-HCl and 100mM NaCl buffer pH 7.5.

### 3. **Figures**

Figure S1. Two-dimensional ^1^H-^15^N HSQC spectrum of LCC^ICCG^-S165A with residue-specific assignment at 50ºC. The reported assignments follow the numbering of crystal structure (PDB code: 6tht) starting at S36 and ending with Q293. The crowded regions are represented in coloured insets. G127 and T211 (black circles) are examples of residues that are broadened near the limit of detection at 50ºC.
